# Rethinking the Estrogen Receptor Beta Dominance Hypothesis in Endometriosis: Insights from Single Cell RNA Sequencing Meta-analysis

**DOI:** 10.1101/2025.09.15.676330

**Authors:** Alexis Heath, Christina Farr Zuend, Wendy A. Goodman, Mehmet Koyuturk, Douglas Brubaker

**Affiliations:** Center for Global Health and Diseases, Department of Pathology, School of Medicine, Case Western Reserve University, Cleveland, Ohio, 44106, USA; Department of Pathology, School of Medicine, Case Western Reserve University, Cleveland, Ohio, 44106, USA; Department of Computer and Data Sciences, Case School of Engineering, Case Western Reserve University, Cleveland, OH, USA; Blood Heart Lung Immunology Research Center of University Hospitals and Case Western Reserve University, Cleveland, Ohio, 44106, USA

**Keywords:** endometriosis, estrogen receptor, beta, alpha, scRNAseq, estrogen, meta-analysis

## Abstract

**Background:**

- Endometriosis is a chronic, estrogen-dependent disease characterized by the presence of endometrial-like tissue growing outside the uterus. The molecular and clinical heterogeneity of endometriosis complicate diagnostic and treatment options -- diagnostic delays of seven to ten years are common and therapies often lack long-term efficacy. Estrogen signaling and estrogen receptor beta (ERβ) expression is thought to be increased in endometriosis, contributing to increased cell proliferation in lesions. The “ERβ dominance hypothesis” is a prevailing hypothesis in the field, setting ERβ as a high-priority therapeutic target. If effectively modulated, ERβ could be the first therapy to directly target lesion biology, rather than only managing symptoms.

**Objective(s):**

- We aimed to characterize ERβ’s expression in endometriosis by cell type and evaluate its therapeutic relevance, primarily assessing the validity of the ERβ dominance hypothesis.

**Study Design:**

- We reanalyzed scRNAseq data from eight previously published studies. Our final filtered dataset included 557,061 cells, the largest endometriosis single cell atlas ever constructed. We quantified gene expression levels of ESR1 and ESR2, which encode ERɑ and ERβ respectively, across each tissue and cell type, to identify cell-type specific drivers of ESR2/ERβ expression across diseased and healthy tissues. To characterize the differences between cells that uniquely express ESR1 versus those that uniquely express ESR2, we performed differential gene expression and pathway enrichment analyses.

**Results:**

- Count and distribution analyses revealed no significant ESR2/ERβ dominance in any cell or tissue type by Fisher’s Exact Tests and Wilcoxon Rank Sum Tests. Differential gene expression and pathway enrichment analyses suggest distinct roles of each estrogen receptor isoform.

**Conclusion(s):**

- Overall, our results argue against a simplified model of ERβ dominance and instead propose a dual-isoform and cell and tissue-specific framework for understanding estrogen receptor signaling in endometriosis. These findings hold important implications for future therapeutic strategies. Specifically, treatments that target ERβ alone may fail to account for the functional role and relative abundance of ERα. In the future, therapeutic approaches that consider isoform-specific, tissue-specific, and cell-specific expression patterns may prove most effective in reducing recurrence and improving outcomes for patients.

**Condensation page:** *Tweetable statement:* Single cell RNA Sequencing meta-analysis shows estrogen receptor beta is not dominantly expressed in most endometriosis tissues. Estrogen receptor alpha to estrogen receptor beta ratios vary by cell type and tissue type. Each isoform directs cell-type specific behavior in endometriosis and disease-free tissues.

*AJOG at a Glance:*

A. Why was this study conducted?
  - We wanted to characterize estrogen receptor beta’s expression in endometriosis and evaluate its therapeutic relevance.
B. What are the key findings?
  - Estrogen receptor beta is not dominantly expressed in any tissue. Estrogen receptor alpha and estrogen receptor beta have disease- and cell-type specific behaviors.
C. What does this study add to what is already known?
  - It characterizes estrogen receptor isoform expression and signaling by cell type. It also challenges the current estrogen receptor beta dominance hypothesis, meaning estrogen receptor beta may not be a key driver of endometriosis.

## INTRODUCTION

Endometriosis is a chronic disease characterized by the presence of endometrial-like tissue growing outside of the uterus that affects approximately 10% of women of reproductive age (1–3). Endometriosis symptoms include dysmenorrhea, chronic pelvic pain, infertility, and a reduced quality of life (4–8), with symptom severity influenced by lesion type, size, depth, and adhesions (9, 10). Molecular alterations in endometriotic lesions such as increased inflammatory mediators, altered hormone signaling, immune dysregulation, and enhanced matrix remodeling are all thought to contribute to lesion development (11–13). The molecular and clinical heterogeneity of endometriosis complicate diagnostic and treatment options -- diagnostic delays of seven to ten years are common, and therapies often lack long-term efficacy (14–19).

Estrogen plays a central role in regulating endometrial tissue growth and function, making its signaling pathway especially relevant in endometriosis. Estrogen signaling is increased in endometriosis and is thought to stimulate cell proliferation in lesions (20–25). With the disease’s established estrogen dependence, prior research has focused on estrogen’s two nuclear receptors -- ERɑ and ERβ -- and their roles in endometriosis pathogenesis (26–35). In the classical estrogen signaling pathway, ERɑ and ERβ bind intracellular estrogens and form dimers which bind to estrogen response elements (EREs) (24). These interactions recruit co-regulatory proteins that can modulate the transcription of estrogen-regulated genes (24).

Increased ERβ expression has been proposed as a key driver of increased estrogen signaling and diminished progesterone responsiveness in endometriotic lesions (36). Prior studies claimed that ERβ is consistently overexpressed in ectopic lesions and ERɑ expression is reduced, shifting lesions to ERβ-dominant signaling (36–41). ERβ is believed to interact with NF-kB to promote inflammatory cytokine production, stimulate COX-2 and PGE2 to increase local estrogen synthesis, and repress ERɑ and progesterone receptor expression to drive progesterone resistance (36). This “ERβ dominance hypothesis” is a prevailing hypothesis in the field, setting ERβ as a high-priority therapeutic target. If effectively modulated, ERβ could be the first therapy to directly target lesion biology, rather than only managing symptoms (42).

Here, we aimed to characterize the prevalence, dominance, and heterogeneity of ERβ’s expression and behavior in endometriosis at the single cell level to better characterize the function of ERβ in endometriosis and provide additional testing of the ERβ dominance hypothesis. We integrated 557,061 cells cross eight filtered unique endometriosis scRNA-sequencing datasets (43–50) to systematically evaluate ERɑ and ERβ expression patterns across multiple tissue types and their associated signaling pathways. Overall, our work provides a comprehensive single-cell-level assessment of whether targeting ERβ could meaningfully impact lesion biology and disease treatment.

## MATERIALS AND METHODS

### Data Preparation and Integration

Data was sourced from NCBI’s Gene Expression Omnibus (GEO) and the Sequence Read Archive (SRA) using keyword searches *“endometriosis/endometrial single cell”* and *“endometriosis/endometrial scRNAseq.”* These search terms produced eight publicly accessible datasets (GSE213216, GSE179640, GSE214411, GSE247695, GSE111976, GSE203191, SRP349751, SRP310621). The data contained cells from thirteen tissue origins (prefix D - disease, C – control) including ectopic lesions (EcE), eutopic endometrium (CEuE and DEuE), endometriomas (EcO), disease-free peritoneal tissue (NED), ovarian tissue (COv and DOv), peritoneal fluid (CPF and DPF), whole menstrual effluent (CWME, DWME), and menstrual effluent tissue (CMET, DMET). Data were downloaded as raw FASTQ files or filtered feature-barcode matrices. FASTQ files were aligned using Cell Ranger to produce filtered feature-barcode matrices.

Each filtered feature-barcode matrix generally represented a single sample from a patient. Each sample was filtered to retain cells with less than 10% mitochondrial content as well as tissue-specific gene count thresholds. Samples from the same tissue types were merged, normalized, scaled, and integrated using reciprocal PCA (RPCA) in Seurat v5.2.0 (51–55). Each newly integrated tissue-specific object underwent Uniform Manifold Approximation and Projection (UMAP) dimensional reduction to enable visualization and clustering of the data (Figure 1).

**Figure 1:**
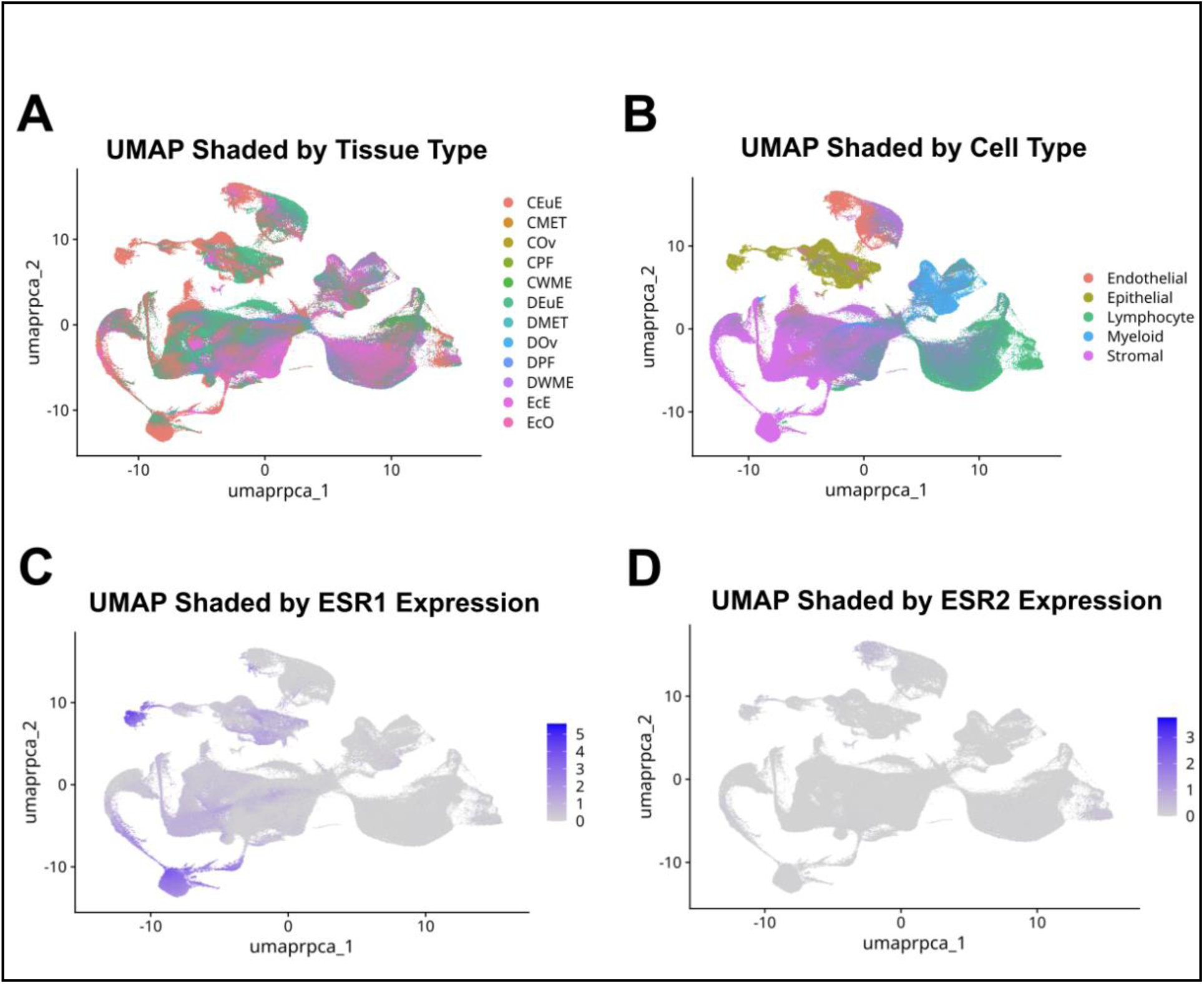
Overall UMAP visual showing 557,061 cells across the thirteen tissue types. Panel A is shaded by tissue origin. Panel B is shaded by assigned cell type. Panel C is shaded by ESR1 Expression. Panel D is shaded by ESR2 Expression.

### Estrogen Receptor Isoform Analysis

We used gene marker lists from the GSE179640 dataset attached in the Supplementary Data to assign cell type identities to each cluster, labeling a cluster as stromal, epithelial, endothelial, myeloid, or lymphocyte. In each cluster, the raw count matrix was extracted to quantify the gene expression levels of ESR1 and ESR2, which encode ERɑ and ERβ respectively. This workflow allowed us to examine the expression patterns of ESR1 and ESR2 in 65 tissue and cell type specific subsets of the data (Supplementary Table 1).

In each tissue and cell type specific subgroup, we compared the distribution of ESR1 expression to ESR2 expression using Wilcoxon Rank Sum tests. We also used Fisher’s Exact Test to examine if ESR1 and ESR2 expression occurred independently of each other on each cell. In addition to these distribution and expression analyses, we calculated the log2(ESR1 expression / ESR2 expression) ratios with a pseudo count of 1 for each cell to further examine relative expression of the isoforms on each cell. We used Wilcoxon Rank Sum testing to compare these ratios across cell types and tissue origins.

### Differential Expression and Pathway Enrichment Analysis

For differential gene expression analysis, cells were partitioned into 4 estrogen receptor isoform groups based on ESR1 and ESR2 expression positivity. Cells were considered uniquely ESR1+ if ESR1 had a count greater than zero and ESR2 expression was zero (Table 1). Cells were considered uniquely ESR2+ if ESR2 had a count greater than zero and ESR1 expression was zero. Cells were considered dually expressing if both ESR1 and ESR2 counts were greater than zero. Cells were considered null expressing if both ESR1 and ESR2 were zero.

**Table 1:** Estrogen receptor isoform expression definitions.

|  | **ESR2 +**<br>(Expression > 0) | **ESR2 –**<br>(Expression = 0) |
| --- | --- | --- |
| **ESR1 +**<br>(Expression > 0) | Dually Expressing | Unique ESR1<br>Expression |
| **ESR1 –**<br>(Expression = 0) | Unique ESR2<br>Expression | Null Expression |

The FindMarkers function was used on each of the 65 subgroups to identify differentially expressed genes (DEGs) in uniquely ESR1+ cells and uniquely ESR2+ cells against null expressing cells. These DEG lists were sorted by descending log2(Fold Change) value and input for Fast Gene Set Enrichment Analysis (FGSEA). These results were filtered to keep pathways with adjusted P values less than 0.05. In each subgroup, the top ten positively enriched pathways and their shared genes were selected for further visualization and comparison.

## RESULTS

### ESR2 Is Not Dominantly Expressed in Any Endometriosis Tissue or Cell Type

Across the eight datasets, we processed 557,061 total cells (210,671 cells from control samples & 346,390 cells from endometriosis samples) for further analysis. These cells came from 134 unique samples – 92 samples from individuals with endometriosis and 42 samples from individuals without endometriosis. Across all five cell types, 148,189 cells were found to be uniquely expressing ESR1. Only 9,889 cells were found to be uniquely expressing ESR2. 5,597 cells expressed both ESR1 and ESR2, and most cells – 393,386 cells – were found to express neither ESR1 nor ESR2 (Supplementary Table 2). To more closely examine expression of the estrogen receptors within and across eutopic and ectopic tissues, disease eutopic tissue (134,100 cells), control eutopic tissue (148,086 cells), ectopic lesions (124,940 cells), and disease-free peritoneum (26,368 cells) were selected as primary tissues of focus for this study.

We used Wilcoxon Rank Sum Tests with Bonferroni correction as a method to identify if the distribution of ESR1 and ESR2 were statistically different within any tissue and cell-type subgroup. There were statistically significant differences in expression distributions between ESR1 and ESR2 in all comparisons with P values ranging from <2.2E-16 to 3.39E-4 (Supplementary Table 3). Contrary to expectations under the ERβ hypothesis, ESR1 visually exhibited overall higher expression across all cell types in our selected tissue types (Supplementary Figure 1).

### Coexpression of ESR1 and ESR2 is the Norm in Endometriosis Tissues and Cell Types

We performed Fisher Exact tests to identify if there was co-regulation or exclusivity of estrogen receptor isoform expression at the cellular level. Across the 20 tissue and cell type specific comparisons in this section, results tended to be tissue-specific (Figure 2). In all control eutopic tissue cell types and disease eutopic tissue cell types, except lymphocytes, results were statistically significant with odds ratios greater than 1, indicating that when one receptor isoform (ESR1 or ESR2) was expressed, the other was significantly statistically likely to be co-expressed rather than a single receptor being expressed the cell type.

**Figure 2:**
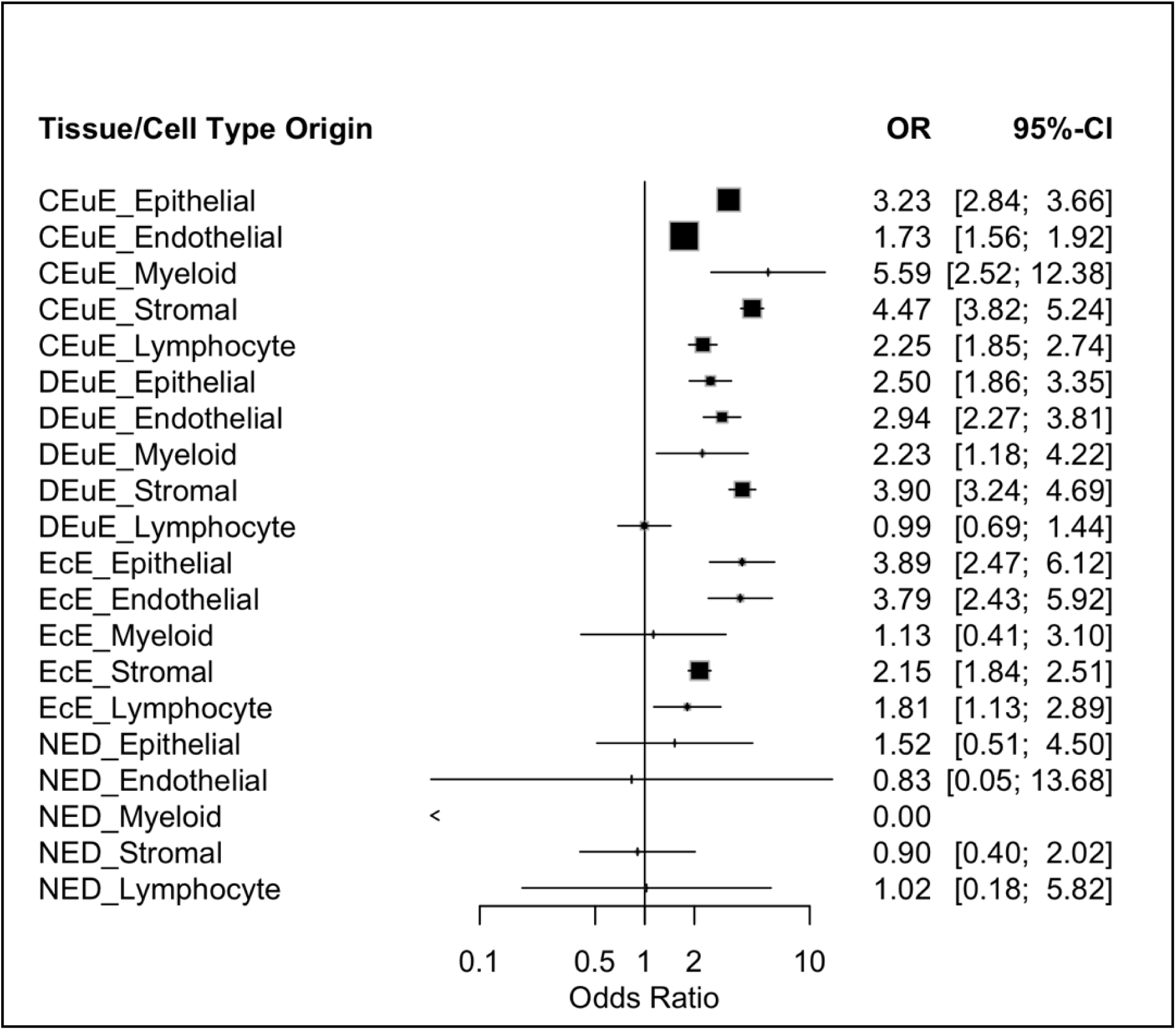
Forest Plot showing odds ratios and confidence intervals from Fisher’s Exact Tests for testing of co-regulation or exclusivity of estrogen receptor isoform expression by tissue and cell type subgroups. Each line represents a tissue (control eutopic endometrium (CEuE), disease eutopic endometrium (DEuE), ectopic endometriotic lesions (EcE), and disease-free peritoneal tissue (NED)) and cell-type (stromal, epithelial, endothelial, myeloid, lymphocyte) specific subgroup. Squares representing the odds ratio are sized based on adjusted p-value with larger squares indicating lower p-values.

The disease eutopic tissue’s lymphocyte cells comparison was not significant (OR = 0.994), showing that ESR1 and ESR2 are not co-expressed and tend to be uniquely and specifically expressed in endometriosis lymphocytes. All ectopic lesion cell types showed significant co-expression except for myeloid cells (OR = 1.126). No disease-free peritoneal tissue cell types were found to be significant, and odds ratios ranged from 0 to 1.5. Therefore, rather than endometriosis cells and tissues being dominated or distinctive in expressing the coding genes for ERβ or ERα, these receptors are most often co-expressed in both eutopic and ectopic tissues.

### The Ratio of ESR1 to ESR2 Expression is Tissue and Cell Type - Specific

Because ESR1 consistently showed higher expression than ESR2 and there was evidence of potential co-regulation, we examined whether the relative expression of ESR1 to ESR2 varied by tissue and cell type. To do so, we compared ESR1/ESR2 expression ratios across cell types and tissue origins using Wilcoxon Rank Sum tests with Bonferroni correction. A statistically significant test would indicate a difference in the continuous distribution of the ESR1/ESR2 ratio between tissues.

When comparing within each tissue type, disease eutopic tissue, ectopic lesions, and disease-free peritoneum, ratio distributions significantly differed across all cell type level comparisons (Supplementary Figure 2). Specifically, the epithelial and stromal cells tended to have more cells with larger positive expression ratios and are explicitly visualized in Figure 3. In control eutopic tissue, all cell type comparisons were significantly different except the epithelial to stromal comparison (Supplementary Figure 2).

**Figure 3:**
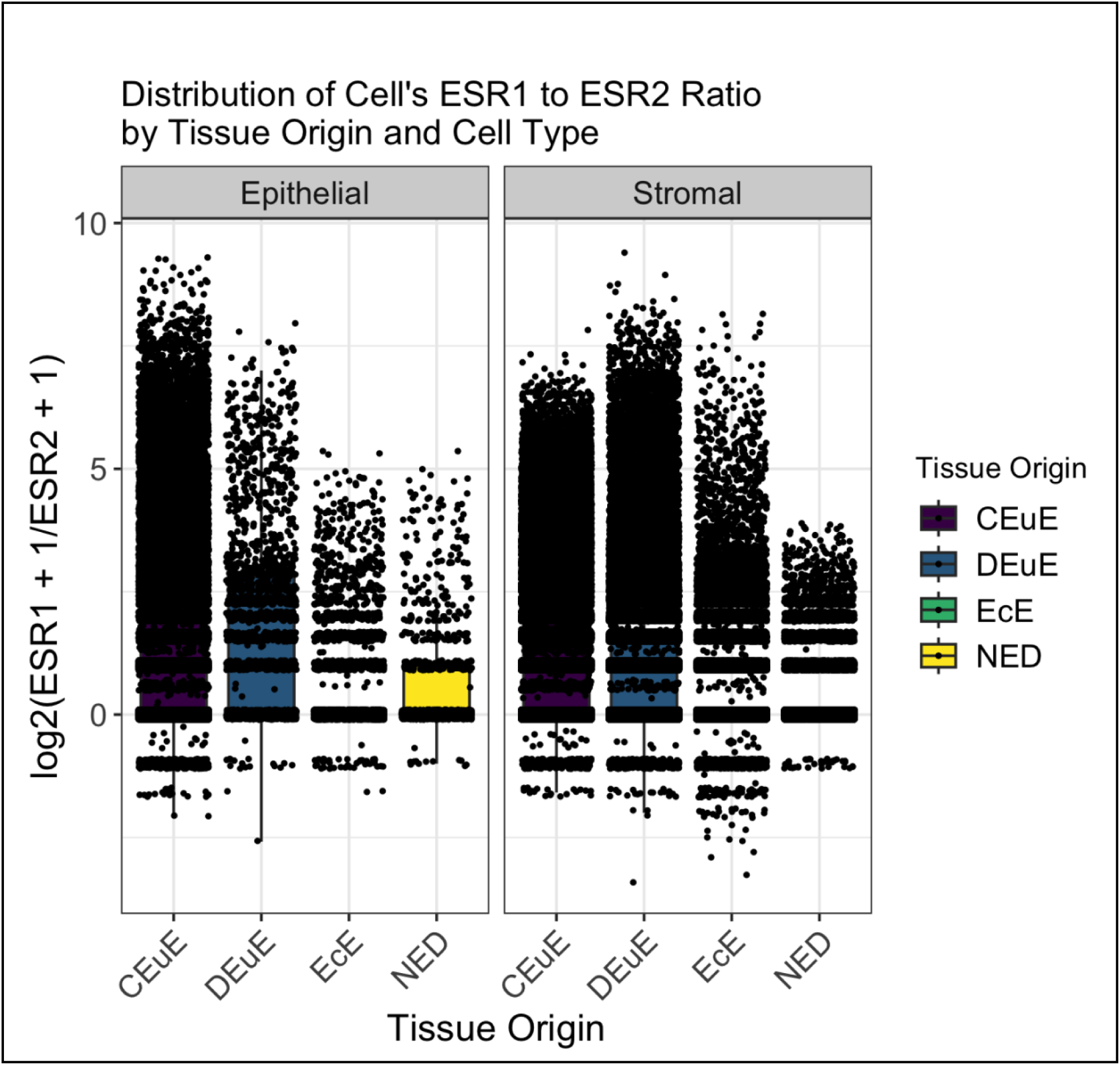
Distribution of single cell ESR1 to ESR2 expression ratios across tissue origins and cell types. Each point represents a single cell’s log2(ESR1 +1 /ESR2 +1) ratio, with boxplots summarizing the distributions in each tissue split by epithelial (left) and stromal (right) cell type. Colors indicate tissue origin: control eutopic endometrium (CEuE), disease eutopic endometrium (DEuE), ectopic endometriotic lesions (EcE), and disease-free peritoneal tissue (NED). A pseudocount of 1 was added to both genes prior to ratio and log transformation. Comparisons between tissue types were made using Wilcoxon Rank Sum tests.

When comparing these distributions by cell type, all comparisons were found to be significantly different except the comparison of disease-free peritoneum to ectopic lesions lymphocyte cells (p value = 0.2) and control eutopic tissue to disease-free peritoneal endothelial cells (p value = 1) (Supplementary Figure 3). Epithelial cells have significantly different distributions in each tissue with eutopic tissues having more larger positive expression ratios compared to ectopic lesions and disease-free peritoneum (Figure 3). Stromal cells were also all significantly different across tissues and appeared visually like the epithelial cells (Figure 3).

Because absolute and relative expression patterns of ESR1 and ESR2 differ across tissues and cell types, we shifted our focus to understand whether these differences could lead to changes in a cell’s downstream functions. In our four selected tissues (CEuE, DEuE, EcE, and NED), we focused on differences between stromal cells exclusively expressing ESR1 or exclusively expressing ESR2 compared to stromal cells expressing neither receptor. This allowed us to identify specific isoform-driven patterns of behavior in the stromal cells of endometriosis and disease-free tissues.

### HLA Genes are Downregulated in ESR1+ Cells from Disease Tissue

ESR1+ stromal cells from control eutopic tissue were positively enriched for multiple pathway types including immune-related pathways, cell signaling pathways, and metabolism pathways. The enrichments were driven by coordinated biological processes, as evidenced by eight of the top ten pathways sharing one or more genes with at least one other pathway (Figure 4A). Of the most commonly occurring shared genes in the ESR1+ stromal cell pathways, all genes were related to the PI3K/Akt and MAPK pathways (Figure 4B). Their coordinated upregulation in ESR1+ stromal cells aligns with the established role of estrogen receptor signaling activating the PI3K/Akt and MAPK pathways to promote proliferation, survival, and tissue remodeling in healthy endometrium. However, these same patterns are not observed among the top ten pathways of DEuE and EcE tissues.

**Figure 4:**
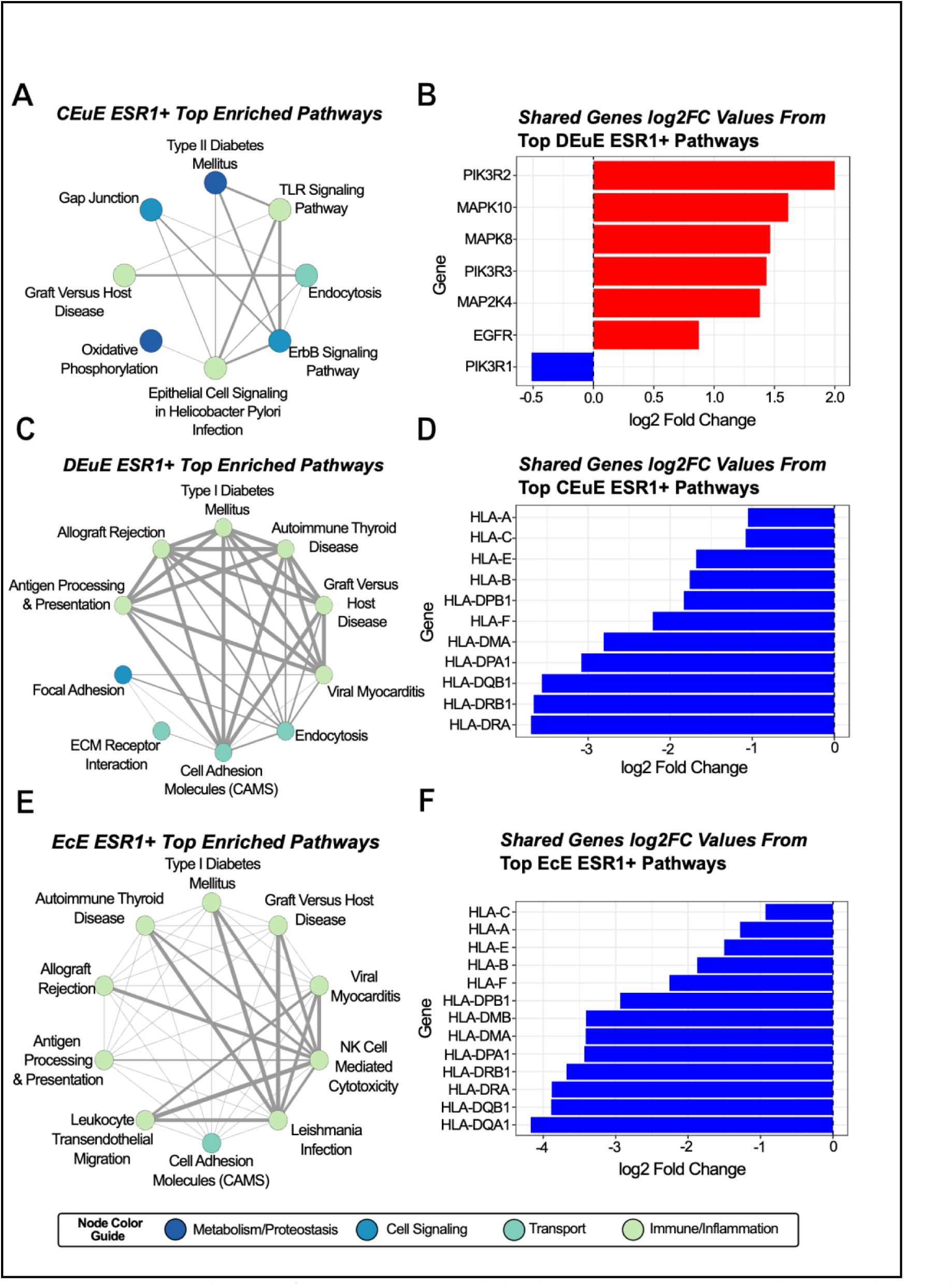
(A-C) Network diagrams of the top positively enriched pathways in each ESR1+ tissue subgroup. Nodes are colored by their main pathway function/group and edge weights are based on the number of shared leading edge genes between pathways, with larger edges indicating more shared genes. (D-F) Bar plots showing the log2(Fold Change) values of the most commonly shared leading edge genes across the top enriched pathways in a ESR1+ tissue subgroup. Red bars indicate a positive log2(Fold Change) while blue bars indicate a negative log2(Fold Change).

In the disease eutopic tissue and ectopic lesion tissues, ESR1+ stromal cells are predominantly enriched for immune-related pathways (Figure 4C-D). Across all these pathways, HLA genes are most shared, and all have significantly negative differential expression values (Figure 4E-F). This indicates diminished antigen-presentation potential in the ESR1+ stromal cells of disease tissue and may contribute to altered immune interactions in disease tissue. ESR1+ stromal cells from disease-free peritoneal tissue had no significant positively enriched pathways.

### MAPK and PI3K/Akt Genes are Enriched in ESR2+ Cells from Control Eutopic Tissue, Disease Eutopic Tissue, and Ectopic Lesions

ESR2+ stromal cells from control eutopic tissue were positively enriched for multiple pathway types including cell signaling pathways, metabolism, and cancer-related pathways (Figure 5A). The most common shared genes from these pathways were involved with the MAPK signaling pathway (Figure 5B). Similar to the ESR1+ stromal cells from the control eutopic tissue, this highlights the central role of MAPK signaling regulated by estrogen receptor signaling in healthy endometrium. However, unlike the ESR1+ stromal cells, the ESR2+ cells in disease eutopic tissue and ectopic lesion tissues shared many of the same enriched genes.

**Figure 5:**
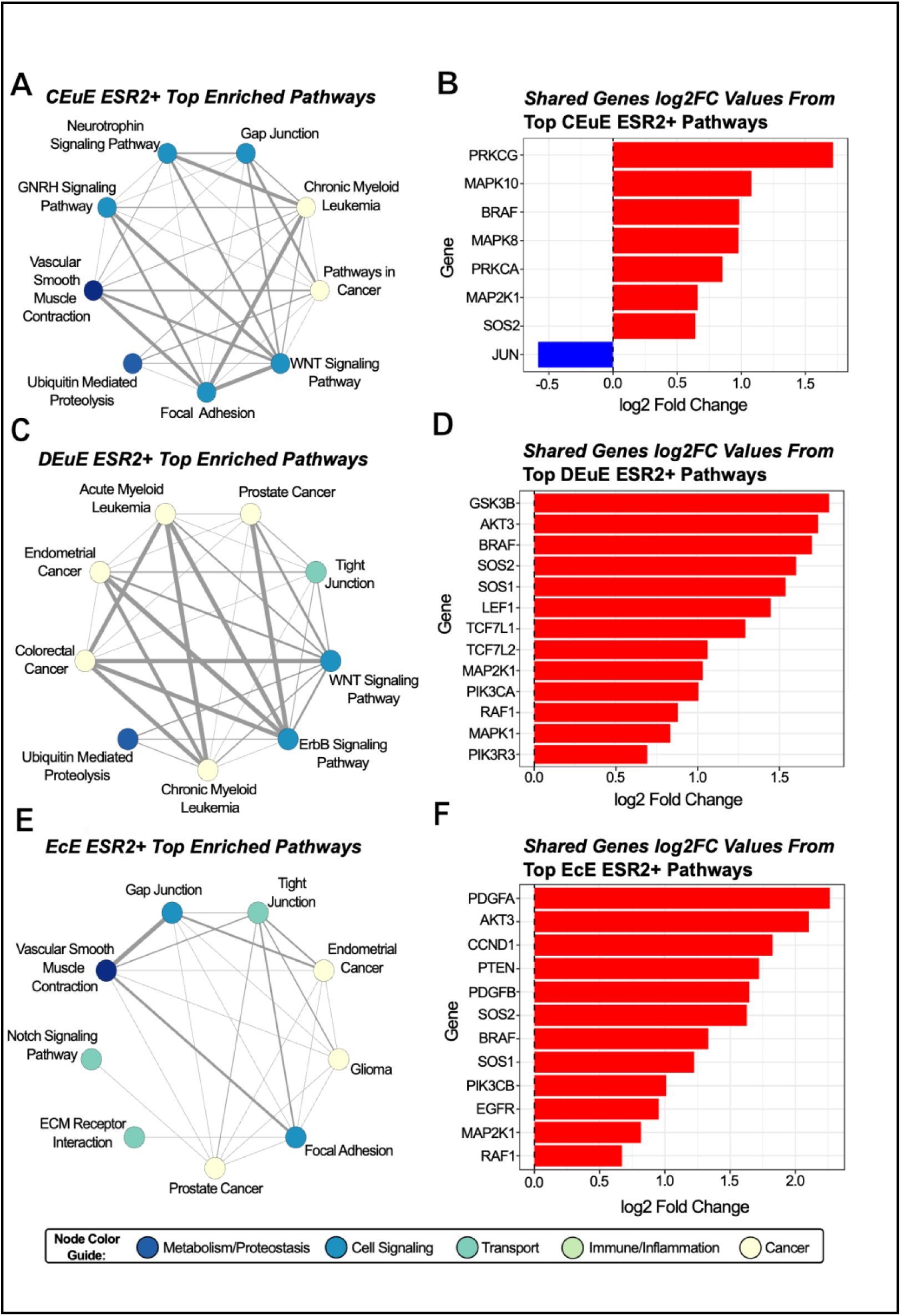
(A-C) Network diagrams of the top positively enriched pathways in each tissue subgroup for ESR2+ cells. Nodes are colored by their main pathway function/group and edge weights are based on the number of shared leading edge genes between pathways, with larger edges indicating more shared genes. (D-F) bar plots showing the log2(Fold Change) values of the most commonly shared leading edge genes across the top enriched pathways in a ESR2+ tissue subgroup. Red bars indicate a positive log2(Fold Change) while blue bars indicate a negative log2(Fold Change). (G-I) Simplified heatmaps for each ESR2+ tissue subgroup showing the normalized enrichment scores for all significantly enriched pathways. An increasingly darker red color indicates a larger positive enrichment score, with values above 2 likely to be biologically meaningful; an increasingly darker blue color indicates a larger negative enrichment score, with values below -2 likely to be biologically meaningful.

ESR2+ stromal cells from disease eutopic tissue and ectopic lesion tissues are predominately enriched in cancer-related pathways (Figure 5C-D). These pathways still most commonly share genes involved in the MAPK pathways in addition to the PI3K/Akt pathway (Figure 5E-F). The ESR2+ stromal cells from disease-free peritoneal tissue had no significantly enriched pathways.

### HLA Gene Expression is Lower in ESR1+ Stromal Cells

To further investigate the expression patterns of the key shared genes driving the pathway enrichments, we examined the average gene expression values of the top shared genes from ESR1+ and ESR2+ stromal cells across control eutopic, disease eutopic and ectopic lesion tissues (Figure 6). This analysis confirmed that HLA genes, which were notably downregulated in the ESR1+ stromal cells from disease tissues are similarly downregulated in the ESR1+ stromal cells from control tissues. However, we do see the HLA genes maintain higher expression in the ESR2+ cells. The top genes involved in the MAPK and PIK3K pathways from the ESR2+ specific analyses had varying expression across the ESR1+ cells. Despite some similar gene expression patterns, these results reinforce isoform- and tissue-specific functional differences observed in the pathway enrichment analyses.

**Figure 6:**
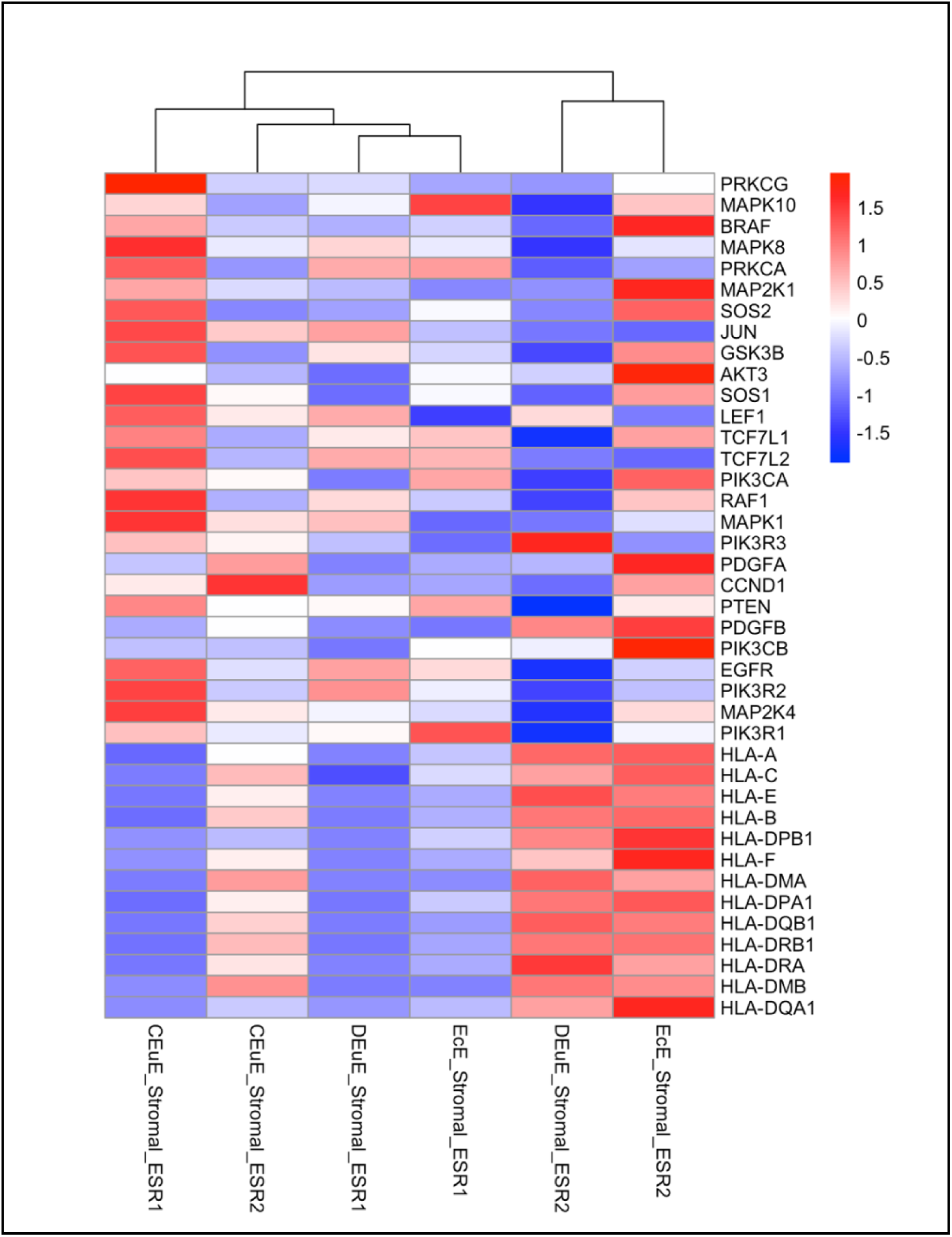
Heatmap of average gene expression values of the top shared leading edge genes from the pathway enrichment analysis. Shared genes from the ESR2+ analysis are at the top of the gene list, followed by four genes seen in both ESR2+ and ESR1+ lists, and ESR1+ specific genes are below that. The gene expression values are log2 transformed and z-scored for easier visualization across a gene. The color gradient means that darker red color indicates higher average expression of that gene in that tissue- and isoform-specific subgroup.

## COMMENT

### Principal Findings

Our analysis of 557,061 cells across eight unique scRNAseq datasets from endometriosis patients reveals no consistent overexpression of ESR2/ERβ in any cell or tissue type. Instead, we observe that the relative expression of ESR1 to ESR2 is both tissue and cell type dependent. ESR1+ stromal cells from control eutopic endometrium have enrichment of pathways and genes linked to PI3K/Akt and MAPK signaling, consistent with known roles of ERα in regulating proliferative and remodeling activities. In disease tissues, we find that the ESR1+ stromal cells shift towards a more prominent enrichment of immune-related pathways with coordinated downregulation of HLA genes. ESR2+ stromal cells also show shared MAPK gene activity across all tissues.

### Results in the Context of What is Known

Prior evidence for ERβ dominance came from bulk tissue mRNA. The ERβ dominance hypothesis relied neither on proteomic evidence nor single cell approaches. Therefore, our test of the hypothesis with scRNA-seq added valuable resolution to the question while also meeting the hypothesis on the terms by which it was established in the field. Our scRNAseq meta-analysis provides a new perspective, showing minimal expression of ERβ and that ERα and ERβ act in complementary roles. Importantly, these roles appear to differ not only by isoform but also by cellular and tissue context.

Through differential gene expression and pathway enrichment analysis, we further demonstrate that cells engage in isoform-specific and disease-specific biological programs. Specific to stromal cells: ESR1+ cells from control eutopic endometrium have enrichment of pathways and genes linked to PI3K/Akt and MAPK signaling, consistent with known roles of ERα in regulating proliferative and remodeling activities. However, in disease tissues, the ESR1+ cells shift towards a more prominent enrichment of immune-related pathways with coordinated downregulation of HLA genes. Though HLA downregulation can also be observed in control ESR1+ cells, the HLA genes are only featured across the top enriched pathways found in disease. The coordinated suppression of these genes could contribute to reduced antigen presentation and potential immune evasion of ESR1+ cells in a disease state.

ESR2+ stromal cells also show shared MAPK gene activity across all tissues. However, because ERβ expression levels are generally much lower than ERα, the overall impact on cellular behavior may be more limited. Together, these findings highlight distinct but complementary roles of ESR1 and ESR2 that vary with disease state.

### Clinical Implications

Our results have important clinical implications for both disease pathogenesis and therapeutic development. If ERβ is not dominant, as observed in our study, ERβ may be less of a key driver than previously thought and ERβ-selective modulators may not be as effective as anticipated. Rather than ERβ-specific therapeutics, therapeutics tailored to ERα, specific tissues, and cell types are likely to be more effective. Most current treatments for endometriosis broadly suppress estrogen signaling, providing symptom relief but offering neither curative nor preventative benefits, and their efficacy often diminishes over time. By characterizing the cell type- and isoform-specific behavior of endometriosis cells like in this study, we can begin designing drugs or estrogen receptor modulators that more precisely target relevant disease-specific pathways.

In addition to estrogen receptor isoform expression differences, we observed coordinated downregulation of HLA genes in ESR1+ stromal cells, particularly in the disease tissues, highlighting a potential link between ERα signaling and immune evasion. This suggests that lesion persistence may result from both aberrant MAPK and PI3K/Akt signaling in estrogen receptor-expressing cells and the ability of these cells to escape immune recognition. Therefore, an effective therapy may need to simultaneously target isoform-specific estrogen receptor signaling and immune evasion mechanisms, allowing for more precise and disease-modifying interventions.

### Research Implications

Protein-level validation of ERα and ERβ expression in stromal and other cell types is essential to confirm the presence and functional relevance of the estrogen receptors as mRNA levels do not always reflect protein abundance or activity. Additionally, functional validation of the isoform-specific signaling pathways, particularly the MAPK/PI3K/Akt activity in ESR1+ and ESR2+ stromal cells, are needed to determine their roles in endometriosis pathophysiology. The coordinated downregulation of HLA genes in ESR1+ stromal cells warrants further investigation to understand potential mechanisms of immune evasion and interactions with the local immune microenvironment. Overall, integrative approaches combing our single-cell transcriptomics results with proteomics or other functional assays will be crucial to fully characterize estrogen receptor isoform roles in specific cell types and tissue contexts.

### Strengths and Limitations

Our study is strengthened by the large number of cells integrated across eight unique datasets. We used appropriate and powered statistical tests to identify differences in count ratio distributions. We performed differential gene expression analyses accounting for the scarcity of single cell RNA sequencing data and held stringent significance thresholds for our pathway enrichment analyses to reduce possible false positive enrichments.

However, there are some limitations to our study. The observational nature of the data limits our ability to establish causal relationships between receptor expression and downstream pathway activation. Further studies would be necessary for functional validation of the top identified pathways and their roles. Additionally, factors such as menstrual cycle stage, exogenous hormone usage, and general patient heterogeneity could influence gene expression patterns and could not be completely controlled for this analysis. Because protein expression and function can be regulated independently of mRNA levels, integrating protein-level analyses in future work with be crucial to fully understanding the biological impact of estrogen receptor isoforms in endometriosis.

### Conclusions

Overall, our results argue against a simplified model of ERβ dominance and instead propose a dual-isoform and cell and tissue-specific framework for understanding estrogen receptor signaling in endometriosis. These findings hold important implications for future therapeutic strategies. Specifically, treatments that target ERβ alone may fail to account for the functional role and relative abundance of ERα. In the future, therapeutic approaches that consider isoform-specific, tissue-specific, and cell-specific expression patterns may prove most effective in reducing recurrence and improving outcomes for patients.

## Acknowledgements

This work was supported by startup funding from Case Western Reserve University, University Hospitals, and the Eunice Kennedy Shriver National Institute of Child Health & Human Development of the National Institutes of Health under Award Number R01HD110367 (to CFZ and DKB). This project was financially assisted by support from The Patty Brisben Foundation for Women’s Sexual Health to CFZ. The views expressed herein do not necessarily represent those of The Patty Brisben Foundation for Women’s Sexual Health.

## Author Contributions

AH: Conceptualization, data curation, formal analysis, investigation, methodology, visualization, writing-original draft, writing-review & editing. CFZ: Conceptualization, funding acquisition, methodology, writing-review & editing. WG: Conceptualization, methodology, writing-review & editing. MK: Conceptualization, methodology, writing-review & editing. DB: Conceptualization, funding acquisition, methodology, project administration, resources, writing review & editing.

## Supplementary Material & Code Availability

All supplementary figures, documents, and code for this analysis are made publicly available at https://github.com/Brubaker-Lab/Characterizing-Estrogen-Receptor-Beta

---

**Supplementary Table 1:**
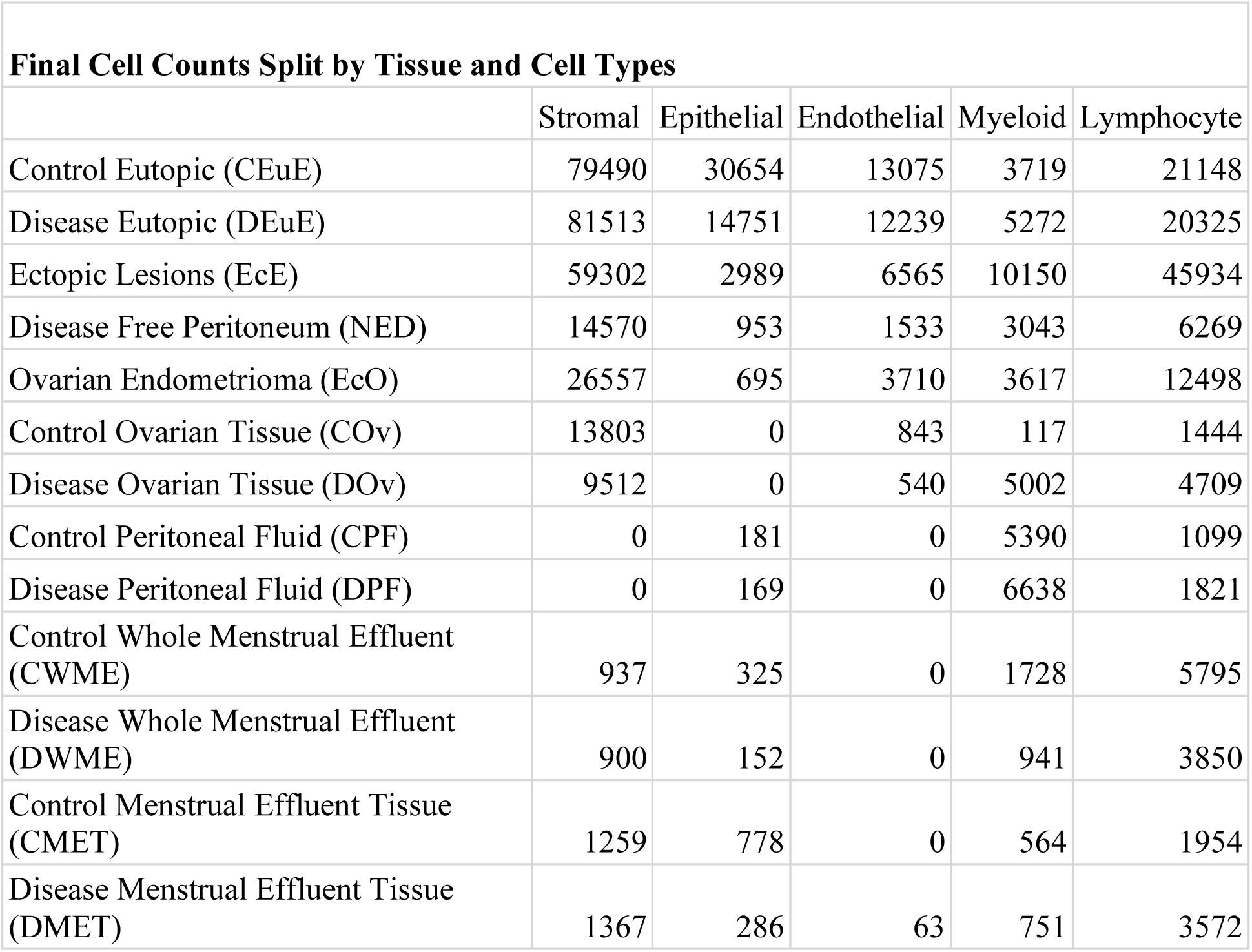
Final filtered cell counts derived from the count matrix for each tissue and cell type subgroup.

| **Final Cell Counts Split by Tissue and Cell Types** |  |  |  |  |  |
| --- | --- | --- | --- | --- | --- |
|  | Stromal | Epithelial | Endothelial | Myeloid | Lymphocyte |
| Control Eutopic (CEuE) | 79490 | 30654 | 13075 | 3719 | 21148 |
| Disease Eutopic (DEuE) | 81513 | 14751 | 12239 | 5272 | 20325 |
| Ectopic Lesions (EcE) | 59302 | 2989 | 6565 | 10150 | 45934 |
| Disease Free Peritoneum (NED) | 14570 | 953 | 1533 | 3043 | 6269 |
| Ovarian Endometrioma (EcO) | 26557 | 695 | 3710 | 3617 | 12498 |
| Control Ovarian Tissue (COv) | 13803 | 0 | 843 | 117 | 1444 |
| Disease Ovarian Tissue (DOv) | 9512 | 0 | 540 | 5002 | 4709 |
| Control Peritoneal Fluid (CPF) | 0 | 181 | 0 | 5390 | 1099 |
| Disease Peritoneal Fluid (DPF) | 0 | 169 | 0 | 6638 | 1821 |
| Control Whole Menstrual Effluent (CWME) | 937 | 325 | 0 | 1728 | 5795 |
| Disease Whole Menstrual Effluent (DWME) | 900 | 152 | 0 | 941 | 3850 |
| Control Menstrual Effluent Tissue (CMET) | 1259 | 778 | 0 | 564 | 1954 |
| Disease Menstrual Effluent Tissue (DMET) | 1367 | 286 | 63 | 751 | 3572 |

**Supplementary Table 2:** Final filtered cell counts derived from the count matrix split by estrogen receptor isoform expression.

| **Final Cell Counts Split by Estrogen Receptor Isoform Expression** |  |  |  |  |  |
| --- | --- | --- | --- | --- | --- |
|  | ESR1+ | ESR2+ | Both | Null | Total |
| Control Eutopic (CEuE) | 73210 | 3199 | 4006 | 67671 | 148086 |
| Disease Eutopic (DEuE) | 52232 | 1048 | 1015 | 79805 | 134100 |
| Ectopic Lesions (EcE) | 12373 | 2699 | 364 | 109504 | 124940 |
| Disease Free Peritoneum (NED) | 3606 | 292 | 19 | 22451 | 26368 |
| Ovarian Endometrioma (EcO) | 2811 | 2126 | 161 | 41979 | 47077 |
| Control Ovarian Tissue (COv) | 827 | 55 | 3 | 15322 | 16207 |
| Disease Ovarian Tissue (DOv) | 372 | 153 | 10 | 19228 | 19763 |
| Control Peritoneal Fluid (CPF) | 916 | 75 | 8 | 5671 | 6670 |
| Disease Peritoneal Fluid (DPF) | 865 | 53 | 8 | 7702 | 8628 |
| Control Whole Menstrual Effluent (CWME) | 293 | 42 | 1 | 8449 | 8785 |
| Disease Whole Menstrual Effluent (DWME) | 124 | 27 | 0 | 5692 | 5843 |
| Control Menstrual Effluent Tissue (CMET) | 213 | 28 | 1 | 4313 | 4555 |
| Disease Menstrual Effluent Tissue (DMET) | 347 | 92 | 1 | 5599 | 6039 |
| Total: | 148189 | 9889 | 5597 | 393386 | 557061 |

**Supplementary Table 3:** All p-values from Wilcoxon Rank Sum tests. We compared the distribution of raw counts of ESR1 to the distribution of raw counts of ESR2. A significant p value indicating that the two distributions are likely to be significantly different from one another. In most cases, ESR1 appeared to visually have higher counts than ESR2, but this did vary, particularly in the EcO tissue.

| **Wilcoxon Rank Sum P Values from ESR1 to ESR2 Comparisons in Each Subgroup** |  |  |  |  |  |
| --- | --- | --- | --- | --- | --- |
|  | Stromal | Epithelial | Endothelial | Myeloid | Lymphocyte |
| Control Eutopic (CEuE) | <2.2e-16 | <2.2e-16 | 5.02e-15 | <2.2e-16 | <2.2e-16 |
| Disease Eutopic (DEuE) | <2.2e-16 | <2.2e-16 | 5.439e-15 | <2.2e-16 | <2.2e-16 |
| Ectopic Lesions (EcE) | <2.2e-16 | <2.2e-16 | <2.2e-16 | 3.672e-10 | <2.2e-16 |
| Disease Free Peritoneum (NED) | <2.2e-16 | <2.2e-16 | 1.656e-11 | 3.667e-15 | 0.0003387 |
| Ovarian Endometrioma (EcO) | <2.2e-16 | 0.2768 | <2.2e-16 | 0.07207 | <2.2e-16 |
| Control Ovarian Tissue (COv) | <2.2e-16 | NA | 0.0178 | 0.3142 | 1.962e-06 |
| Disease Ovarian Tissue (DOv) | 2.128e-09 | NA | 0.1792 | 3.278e-10 | 7.896e-06 |
| Control Peritoneal Fluid (CPF) | NA | 0.5256 | NA | <2.2e-16 | 0.002244 |
| Disease Peritoneal Fluid (DPF) | NA | 0.04713 | NA | <2.2e-16 | 0.02089 |
| Control Whole Menstrual Effluent (CWME) | <2.2e-16 | 0.006243 | NA | 5.816e-11 | 2.25e-05 |
| Disease Whole Menstrual Effluent (DWME) | <2.2e-16 | 0.0001847 | NA | 0.03219 | 0.0001412 |
| Control Menstrual Effluent Tissue (CMET) | <2.2e-16 | 2.825e-11 | NA | 0.02027 | 0.002167 |
| Disease Menstrual Effluent Tissue (DMET) | <2.2e-16 | 2.022e-13 | 0.3251 | 0.002611 | 2.948e-11 |

**Supplementary Figure 1:**
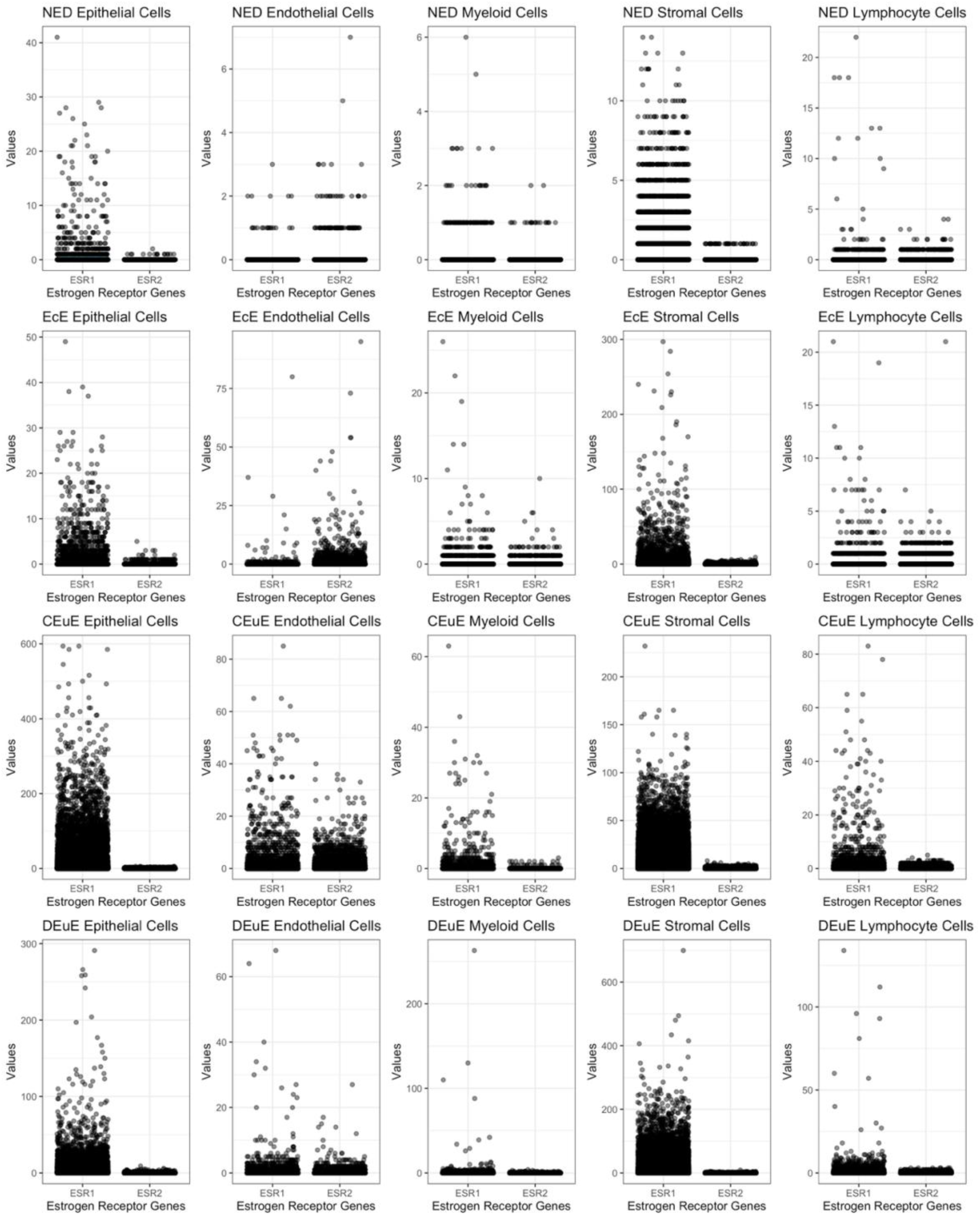
Each boxplot shows the distribution of raw counts for either ESR1 or ESR2. Each panel represents a specific tissue and cell type subset – each row also represents one tissue while each column represents a cell type. ESR1 visually is expressed higher than ESR2 in general.

**Supplementary Figure 2:**
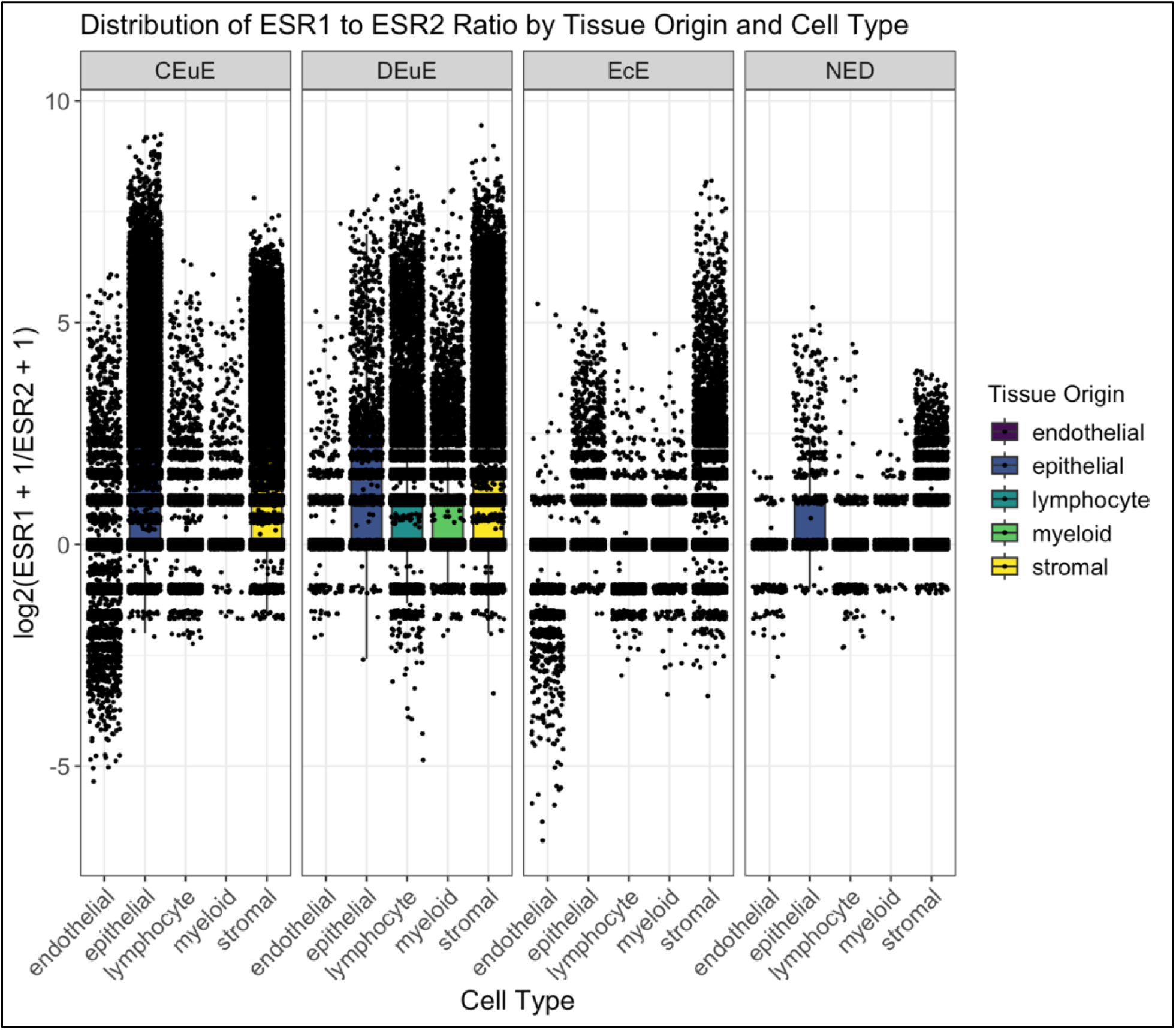
These boxplots are split into four panels, containing one panel for each tissue type – Control Eutopic Endometrium (CEuE), Disease Eutopic Endometrium (DEuE), Ectopic Lesions (EcE), Disease-Free Peritoneum (NED). In each panel, there is a boxplot for each identified cell type containing a point for every cell’s log2(ESR1/ESR2) in that subgroup. When comparing distributions of these ratios using Wilcoxon Rank Sum between cell types in each tissue origin separately, every comparison was significant (p value <0.05) except for one. In CEuE tissue, the epithelial to stromal comparison was insignificant, indicating the distribution of these ratios were not significantly different from one another.

**Supplementary Figure 3:**
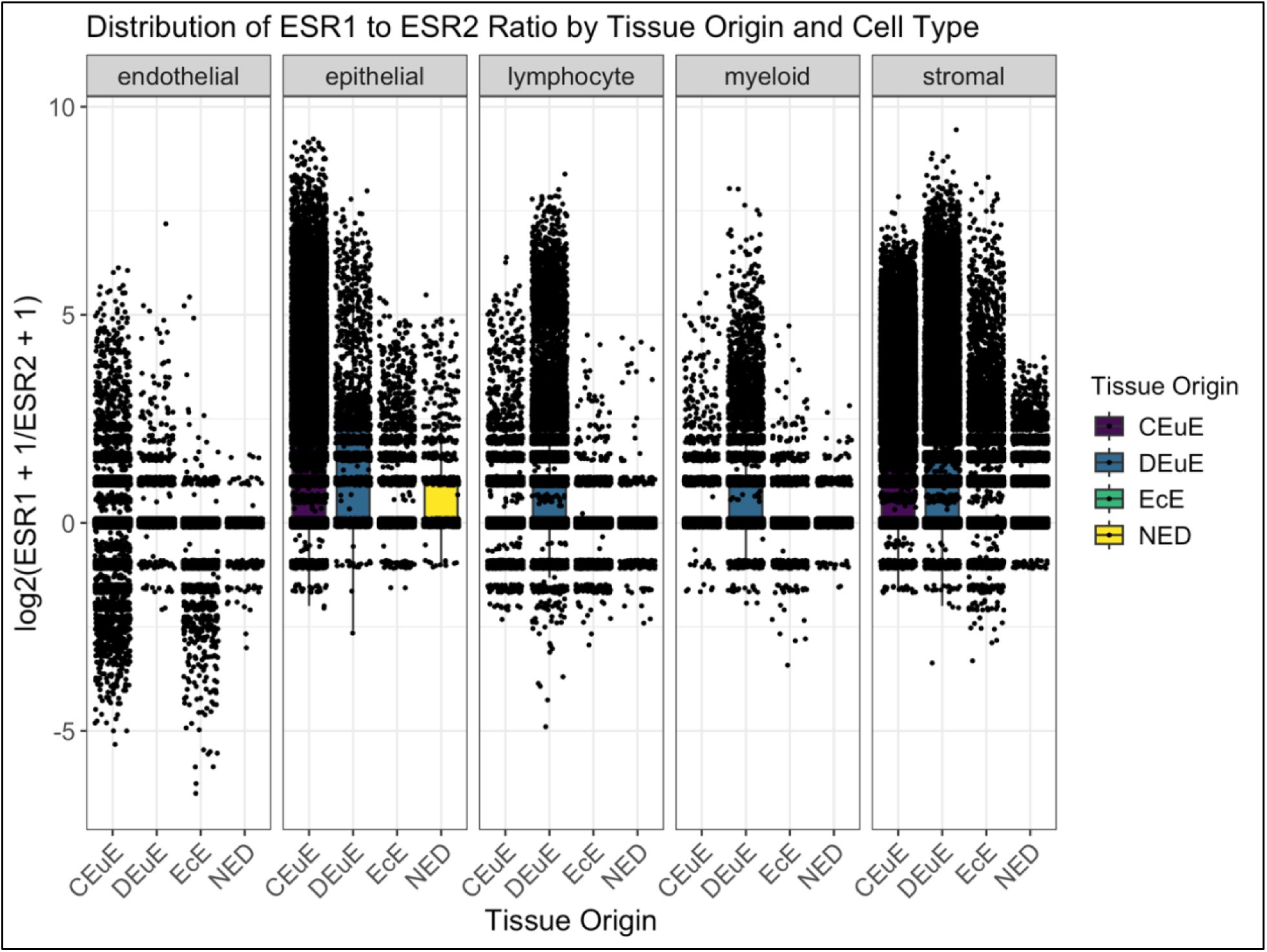
These boxplots are split into five panels, containing one panel for each cell type – endothelial, epithelial, lymphocyte, myeloid, and stromal. In each panel, there is a boxplot for each tissue type – Control Eutopic Endometrium (CEuE), Disease Eutopic Endometrium (DEuE), Ectopic Lesions (EcE), Disease-Free Peritoneum (NED) – containing a point for every cell’s log2(ESR1/ESR2) in that subgroup. When comparing distributions of these ratios using Wilcoxon Rank Sum between tissue origins in each cell type separately, every comparison was significant (p value <0.05) except for two. NED to EcE lymphocyte cells (p value = 0.2) and CEuE to NED endothelial cells (p value = 1).

**Supplementary Figure 4:**
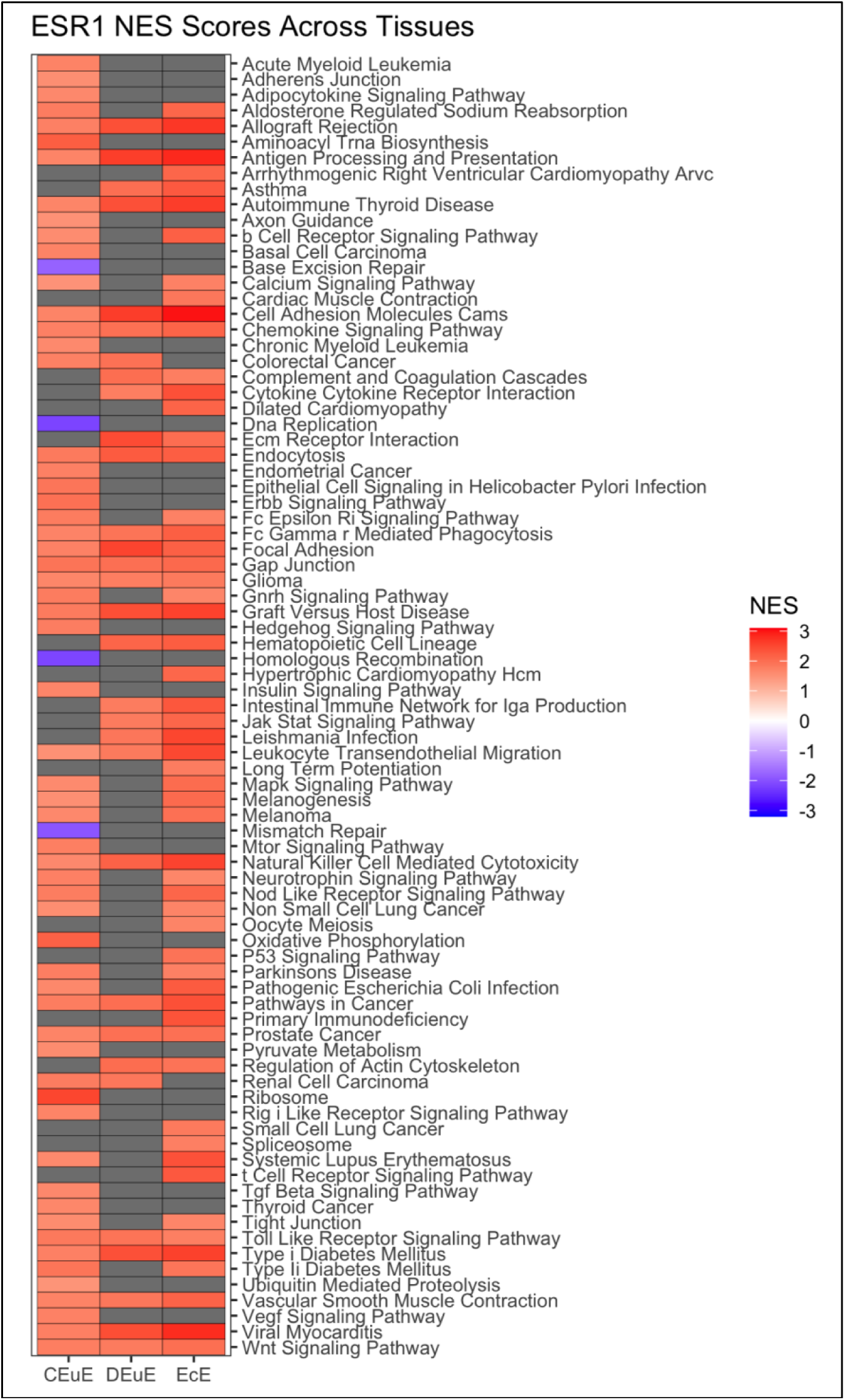
This heatmap compares the normalized enrichment scores of all enriched pathways with adjusted p values less than 0.01 from ESR1+ cells compared to null expressing cells across three tissue types: Control Eutopic Endometrium (CEuE), Disease Eutopic Endometrium (DEuE), and Ectopic Lesions (EcE). The gradient ranges from blue to red with a darker blue indicating a more negative normalized enrichment score and darker red indicating a more positive normalized enrichment score.

**Supplementary Figure 5:**
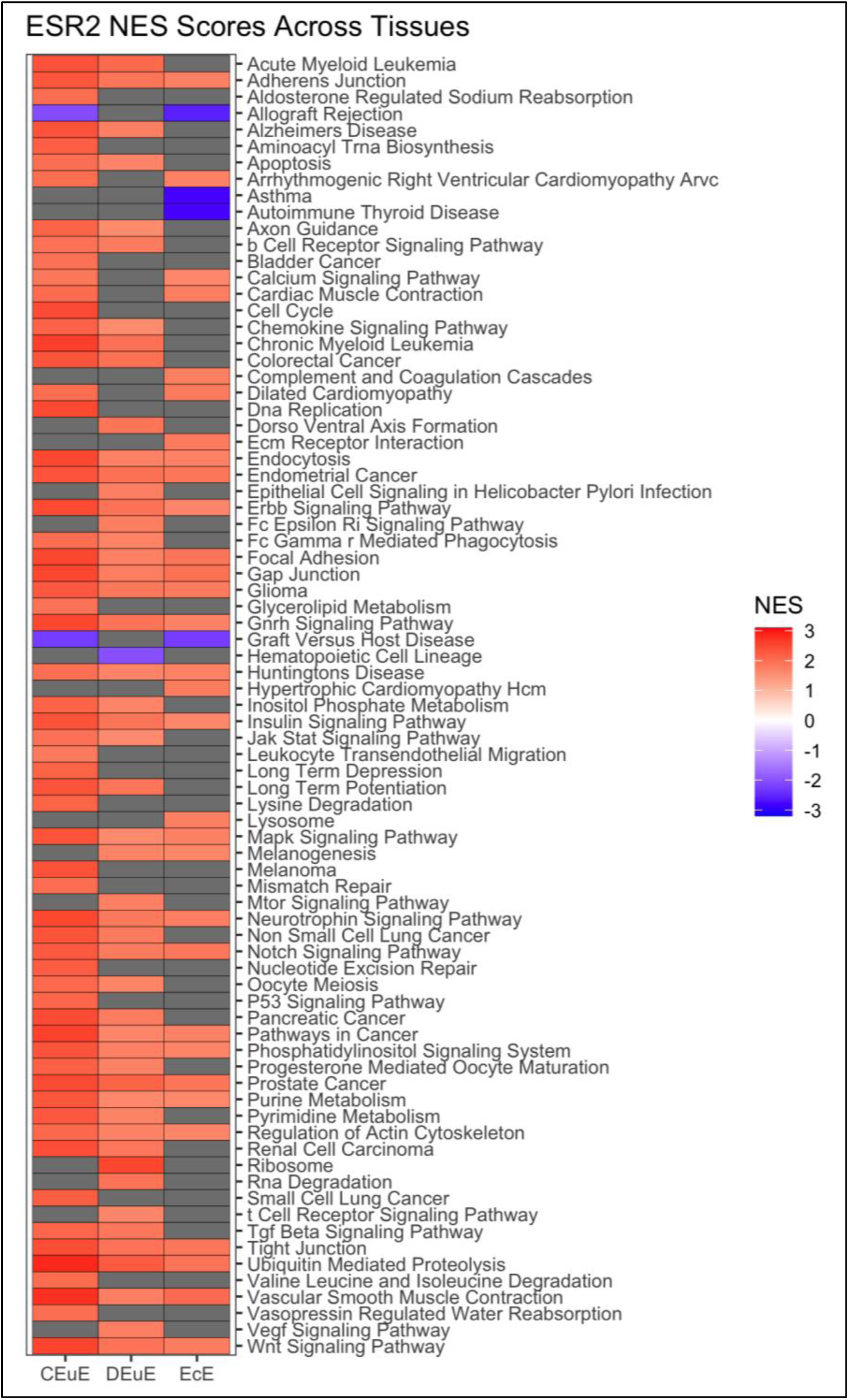
This heatmap compares the normalized enrichment scores of all enriched pathways with adjusted p values less than 0.01 from ESR2+ cells compared to null expressing cells across three tissue types: Control Eutopic Endometrium (CEuE), Disease Eutopic Endometrium (DEuE), and Ectopic Lesions (EcE). The gradient ranges from blue to red with a darker blue indicating a more negative normalized enrichment score and darker red indicating a more positive normalized enrichment score.

**Supplementary Figure 6:**
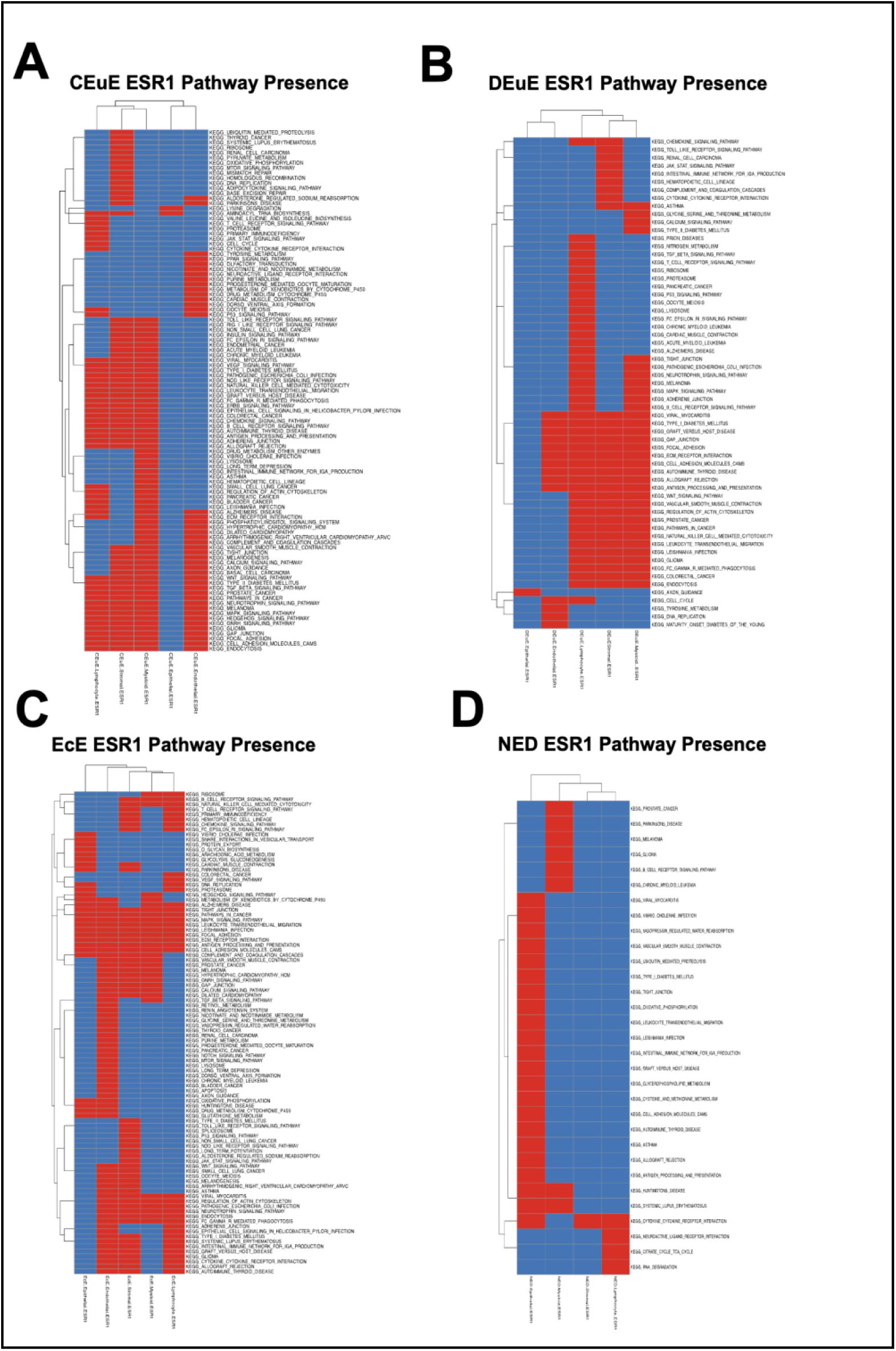
Pathway Presence across ESR1+ cell types in each tissue origin of interest. Red color indicates the pathway is significantly enriched in that subgroup. Blue color indicates the pathway is not significantly enriched in that subgroup.

**Supplementary Figure 7:**
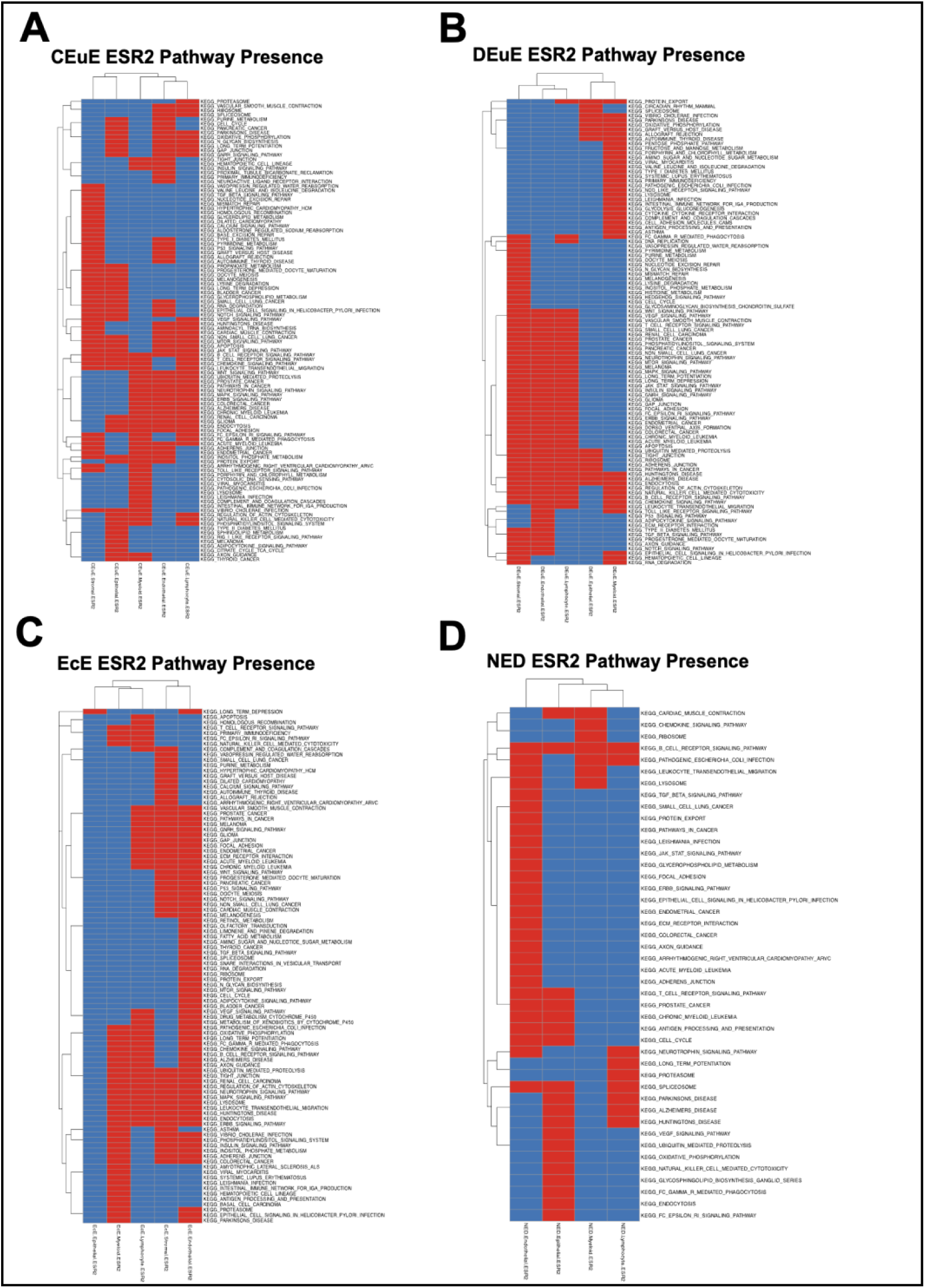
Pathway Presence across ESR2+ cell types in each tissue origin of interest. Red color indicates the pathway is significantly enriched in that subgroup. Blue color indicates the pathway is not significantly enriched in that subgroup.

